# In vivo characterization of the connectivity and subcomponents of the human globus pallidus

**DOI:** 10.1101/017806

**Authors:** Patrick Beukema, Fang-Cheng Yeh, Timothy Verstynen

**Author notes:** Corresponding Author: Carnegie Mellon University Department of Psychology 340 U Baker Hall Pittsburgh, PA 15213 (.

## Abstract

Projections from the substantia nigra and striatum traverse through the pallidum on the way to their targets. To date, in vivo characterization of these pathways remains elusive. Here we used high angular resolution diffusion imaging (N=138) to study the characteristics and structural subcompartments of the human pallidum. Our results show that the diffusion orientation distribution at the pallidum is asymmetrically oriented in a dorsolateral direction, consistent with the orientation of underlying fiber systems. Furthermore, compared to the outer pallidal segment, the internal segment has more peaks in the orientation distribution function and stronger anisotropy in the primary fiber direction, consistent with known cellular differences between the underlying nuclei. These differences in orientation, complexity, and degree of anisotropy are sufficiently robust to automatically segment the pallidal nuclei using diffusion properties. Thus the gray matter diffusion signal can be useful as an in vivo measure of the collective nigrostriatal and striatonigral pathways.

## Introduction

The basal ganglia are a crucial forebrain network associated with many cognitive functions, including reward processing, decision-making, and learning (Hollerman et al., 2000; Haber, 2003). Many aspects of basal ganglia function rely on dopaminergic inputs from the substantia nigra that serve as a modulatory signal for neurons in the subpallium (Haber et al., 2000). These dopaminergic inputs are conducted by a set of fiber bundles that originate in the pars compacta region of the substantia nigra, a portion of which migrate in a dorsolateral direction through the segments of the globus pallidus (Carpenter and Peter, 1972). While a majority of these projections pass through the pallidum and terminate on cells in the striatal nuclei (Carpenter and McMasters, 1964), a significant number of them also terminate in the inner and outer segments of the globus pallidus, forming the nigropallidal pathway (Cossette et al., 1999). Two other major fiber systems traversing through the globus pallidus project from the striatum, including the striatopallidal fiber systems, which form the canonical direct and indirect pathways, and the striatonigral fiber system. Breakdowns in these various pathways form the etiology of several neurodegenerative diseases. For example, axonal degeneration of the nigrostriatal pathway is the pathological hallmark of Parkinson’s Disease (Burke and O’Malley, 2013) while Huntington’s disease is characterized by a loss of medium spiny neurons within the striatum that project to the globus pallidus (Reiner et al., 1988). Thus *in vivo* characterization of these basal ganglia projections has clear clinical implications.

One problem with characterizing both the nigral and striatal efferents is that they are largely embedded within the gray matter of several basal ganglia nuclei, primarily within the globus pallidus. The globus pallidus is the primary output of the basal ganglia network, sending projections that relay signals from upstream nuclei to the thalamus (Alexander et al., 1986). In primates it is comprised of an external segment (GPe), which serves as an inhibitory relay nucleus within the indirect pathway, and the internal segment (GPi) that aggregates all information from all basal ganglia pathways. While primarily defined by their connectivity and neurophysiological profiles, the GPe and GPi are also distinguishable at the cellular level by differences in cell density, cell type, and morphology (Hardman et al., 2002, Eid et al 2013, Difiglia and Rafols, 1988). One of the most salient differences is that the GPi has a much lower overall neuronal density (see table 2 in Hardman et al., 2002). More importantly, given that the volume of the GPi is smaller than that of the GPe and that both nigrostriatal and striatonigral fibers pass through the globus pallidus on their way to their targets, the GPi also has a greater density of both nigrostriatal and striatonigral efferents than its external counterpart.

Despite the clear morphological differences, in vivo characterization of these critical nuclei remains elusive by MRI-based neuroimaging technologies, particularly with conventional T1-weighted or T2-wighted images used in most human neuroimaging experiments. This is because T1-weighted and T2-weighted approaches have limited power to characterize the microscopic structure of the GPe and the GPi. This limitation can be compensated by the recent advances in high angular resolution diffusion MRI, which offers a non-invasive approach to study microscopic structure. One advantage of diffusion MRI is that it is able to detect microscopic differences in underlying cellular morphologies, including spatial asymmetry in underlying axonal tracts, and it has shown great promise in quantifying variability in underlying tissue composition (Behrens et al., 2003). With these unique features, diffusion MRI has emerged as an increasingly popular tool for characterizing the neural tissues (Abhinav et al., 2014). While most commonly used to study structural subcomponents of large white matter fascicles (Bastiani, 2012, Wang et al, 2013, Fernandez-Miranda 2014), diffusion MRI has also been shown to be useful for characterizing differences in microstructural cellular properties of gray matter (Wiegell et al., 2003; Mang et al., 2012), including sensitivity to both neural and glial distribution patterns (Blumenfeld-Katzir et al., 2011). For example, analysis of the diffusion MRI signal has been used to segment the different thalamic nuclei in humans (Behrens et al, 2003).

Here we adopt an atlas approach to study the orientation distribution functions of the water diffusion, termed spin distribution functions (SDF; Yeh et al. 2010) within the gray matter of the globus pallidus in a stereotaxic space. An SDF provides a nonparametric representation of the diffusion pattern that cannot be offered by conventional tensor-based analysis, thus allowing for characterizing and segmenting structural subcomponents. Here we used data from two high angular resolution diffusion sequences, diffusion spectrum imaging (DSI) and multi-shell imaging (MSI) and examined the diffusion characteristics in the nigral and striatal efferents. We also examined whether there are reliable differences in the SDFs between the GPe and the GPi that allows for accurate segmentation based solely on diffusion properties. These findings may identify a clear potential for using high angular resolution diffusion MRI as a novel *in vivo* characterization of the microarchitecture of the human globus pallidus, including the nigral and striatal efferents that break down in various neurological pathologies.

## Results

### Dorsolateral orientation of the pallidal diffusion signal

The nigrostriatal and striatonigral fibers traverse the pallidum on the way to their targets, resulting in a primarily dorsolateral-ventromedial orientation of the collective axons (Fig. 1A). In humans, the two segments of the pallidum are separated by a thin white matter band, called the internal medullary lamina, shown in coronal sections from the Big Brain atlas in Fig.1B and 1C (Amunts et al., 2013). To characterize the orientations of the fibers in the inner and outer segment of the globus pallidus, we isolated the voxels corresponding to the two segments (Methods, Fig 2A). Within each of these region masks we estimated the SDFs from the diffusion signal of each voxel (Fig 2B-D), which is a 3D representation of the underlying diffusion orientation distribution. The first three peaks in each SDF were extracted and their orientation and anisotropy intensity, called quantitative anisotropy (QA; Fig. 2B, see Methods), were recorded for every voxel in each mask. Two representative SDFs from an example subject from the DSI dataset show how the shapes of the SDFs differ between the inner (Fig. 2C) and outer (Fig. 2D) segments of the pallidum.

**Fig. 1.**
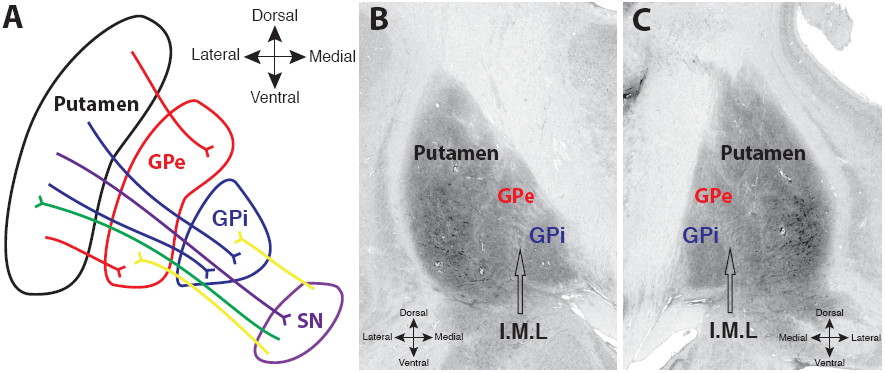
A. Schematic of fiber systems traversing the internal (GPi) and external (GPe) segments of the pallidum, including projections to and from the substantia nigra. Labeled pathways include nigrostriatal (green) striatopallidal (red, cyan), striatalnigral (purple) and nigropallidal (yellow (red, blue), striatonigral (purple) and nigropallidal (yellow). B and C. Coronal images from the Big Brain atlas, showing the putamen, GPe, and GPi, in the left (B) and right (C) hemispheres; black arrows point to the approximate center of the internal medullary lamina (I.M.L) separating the two segments (slice 3894, Amunts 2013).

**Fig. 2.**
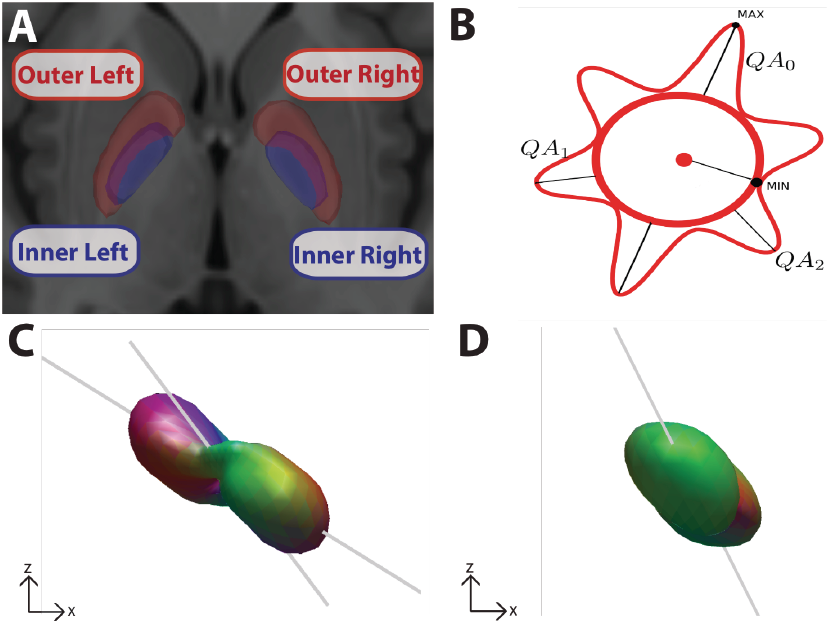
A.The inner (blue) and outer (red) segments of the left and right pallidum were manually drawn on the high resolution T1 ICBM 152 template. B. Schematized version of an SDF illustrating three resolved fibers (QA0, QA1, QA2), their magnitude (i.e., lengths, reflecting QA) and orientation. C,D Representative SDFs from the left internal segment (C) and left external segment (D) in the coronal plane from a single subject from the DSI dataset. Gray lines indicate direction of fiber orientations.

Figure 3 shows the distribution of principal fiber angles across subjects in the DSI and MSI samples. Each distribution was confirmed to be non-uniform using a Rayleigh’s test (z values > 36.12, p-values < 0.001) and exhibited consistent peaks in the orientations in the fibers in both region masks and both data samples. In both the DSI and MSI samples, the distribution of the angles in the inner segment is predominately concentrated in the dorsolateral direction, consistent with the known orientation of the major nigral and striatal efferents (Wilson 1914, Szabo 1967, Szabo 1962, Fox and Rafols, 1976, Carpenter and Peter 1972). Compared to the inner segment, the distribution of the angles in the outer segment is rotated medially, clockwise in the left hemisphere and anticlockwise in the right hemisphere. Interestingly, this rotation is more pronounced in the DSI dataset, resulting in the majority of fibers pointing dorsomedially (towards the internal capsule). One possibility is that the predominantly dorsal and dorsolateral fiber systems are not contributing to the strongest anisotropy pattern in the DSI sample, but are still present at lower anisotropy thresholds. To explore this, we looked at the orientation of the secondary fibers in both datasets (Fig. 4). Indeed, the secondary fiber in the DSI dataset was oriented in a more dorsolateral direction as predicted if it were reflecting the angle of the underlying nigral and striatal efferents (red histogram in Fig. 4A,B).

**Fig. 3.**
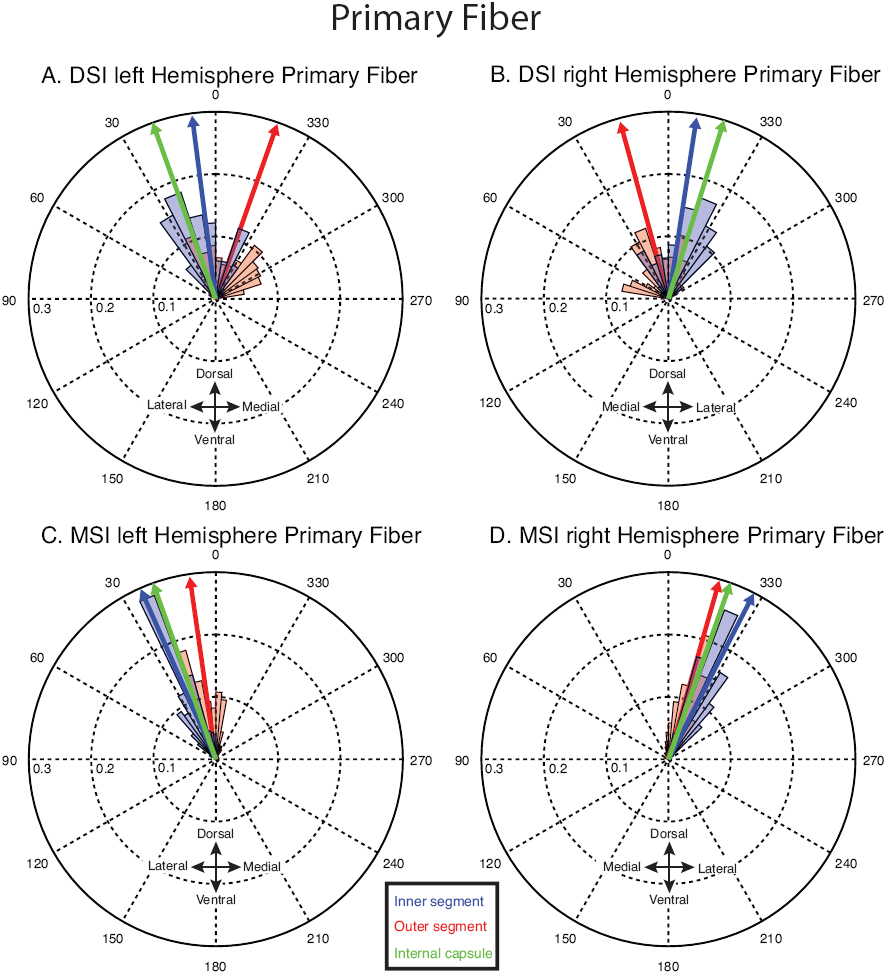
Angular distributions of primary fibers from DSI (A,C) and MSI (C,D) datasets. Blue, red, and green arrows indicate the mean of the distributions in the inner, outer and internal capsule respectively.

**Fig. 4.**
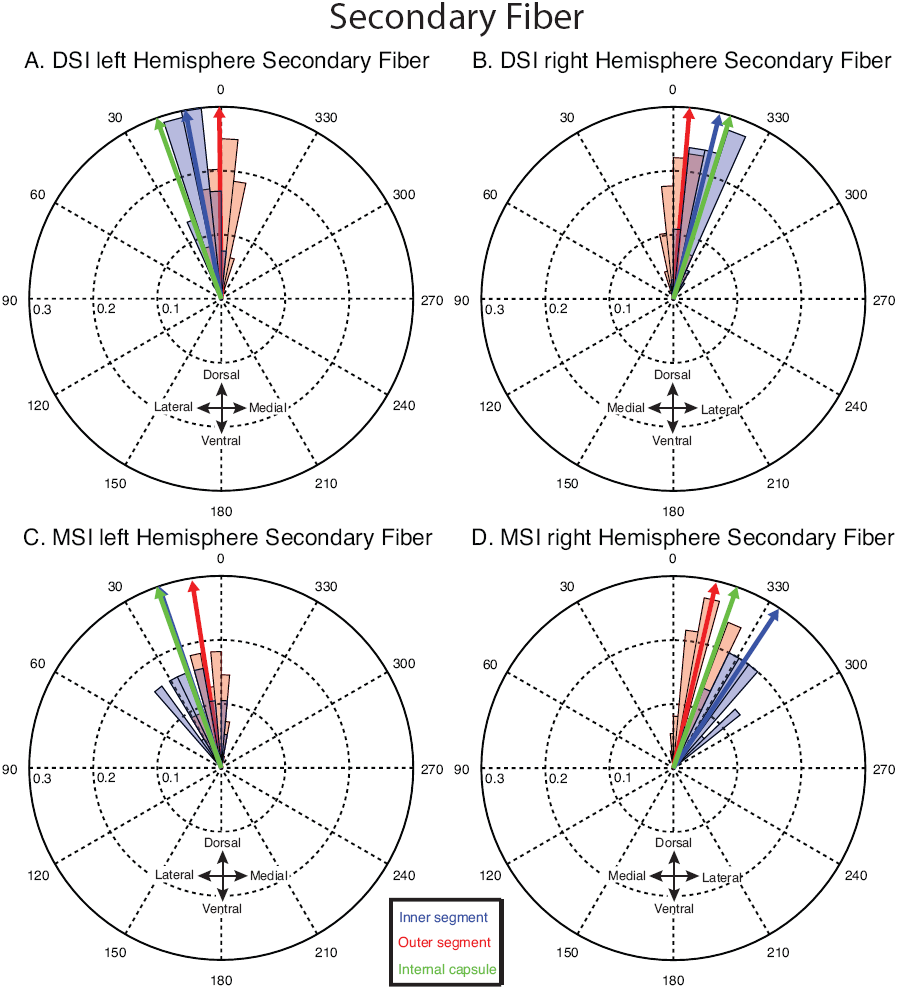
Angular distributions of secondary fibers from DSI (A,C) and MSI (C,D) datasets. Blue, red, and green arrows indicate the mean of the distributions in the inner, outer and internal capsule respectively.

These patterns of fiber peak orientations within the SDF are also clearly visible in the geometries of the extracted fibers at each voxel. Figure 5 shows a coronal slice from two representative subjects from the DSI and MSI samples. A majority of the primary and secondary fibers overlap in the inner segment (blue voxels) and are generally oriented dorsolaterally, whereas the primary and secondary fibers in the external segment (red voxels) show less overlap and exhibit a greater abundance of dorsomedial orientations compared to the internal segment. Thus, the gray matter diffusion signal within the pallidal nuclei has asymmetries in peak anisotropy directions that are consistent with the orientation of nigral and striatal efferents running through the pallidum, suggesting that the diffusion signal is sensitive to these underlying pathways.

**Fig. 5.**
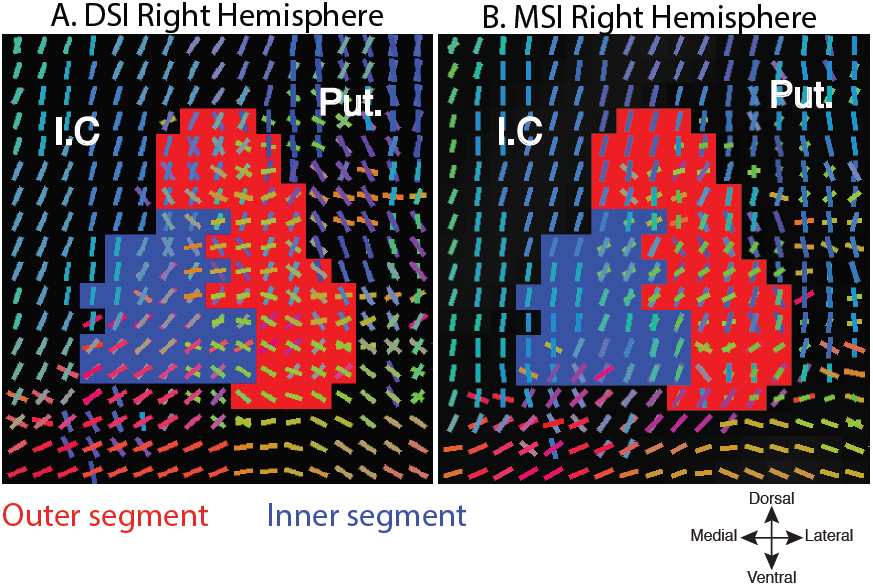
Voxelwise geometries of the primary and secondary fibers, inner (blue) and outer (red), segments of two example subjects from the DSI (A) and MSI (B) datasets. Slices are both from y = -1 (MNI). (Putamen (Put.), Internal capsule (I.C.). Fiber orientations are color coded according to their orientation.

### Differential diffusion patterns between inner & outer pallidal segments

Along with differences in fiber orientation, we also observed general differences in the sensitivity and intensity of the diffusion signal between the two pallidal segments. Tensor-based analyses have shown that the anisotropy patterns around the principal fiber direction tend to be highly sensitive to underlying cellular morphology differences (Wiegell et al., 2003). Figure 6, panels A through D, shows the across-subject probability distributions of the mean QA in the principal fiber direction, for both pallidal segments. The peaks (arrows) of the distributions occurred at consistently higher thresholds in the inner segment in both hemispheres and both samples. Thus, the inner segment exhibited a mean shift in QA compared to the outer segment mask, suggesting a slightly stronger diffusion intensity for the inner pallidal segment.

**Fig. 6.**
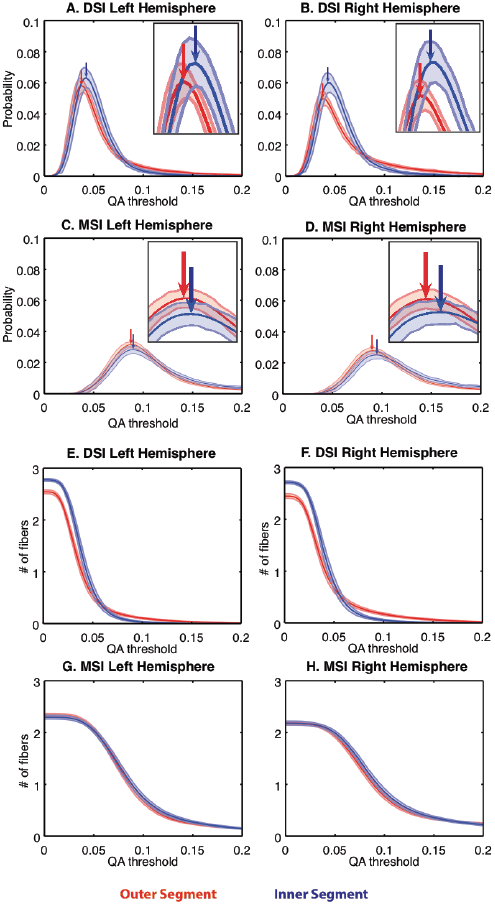
A-D. Probability density functions of the primary fiber QA of the inner (blue) and outer (red) segments in the DSI(A,B) and MSI(C,D) datasets. E-H The number of resolved fibers thresholded by QA in the inner and outer segments in the DSI (A,B) and MSI datasets (C,D). Arrows indicate peaks of the distributions. Lines indicate mean and shaded regions are 95% confidence intervals.

Differences between the pallidal segments were also reflected in the complexity of the SDF geometry. The sensitivity curves in Figure 6, panels E through H, show the average number of resolved fibers (y-axis) with QA values above the value shown on the x-axis. Thus this measures the complexity of the diffusion geometry in each voxel by showing the robustness of the fiber peaks within the reconstructed SDF across a range of thresholds. As expected, the number of resolved fibers decays rapidly as the threshold increases. Notably, compared to the outer segment, in the DSI sample we detected more fibers in the inner segment at thresholds less than QA = 0.05 (Fig.6 E,F), and more fibers in the outer segment at thresholds higher than 0.05. In the MSI sample, these distributions largely overlapped (Fig.6 G,H), although the inner segment in the right hemisphere showed a smaller shift in the same direction as was observed in the DSI dataset. Taken together, the differences in orientation, intensity and sensitivity between the structures suggest that the diffusion signal is picking up on reliable differences in the cellular content of the two nuclei.

### Reliable segmentation of pallidal nuclei

If these differences are reflecting distinctive cellular architectures and local connectivity patterns then it should be possible to classify the two segments based purely on the properties of the DWI signal. To this end we used k-means clustering to segment all voxels within the globus pallidus using three voxel features as inputs: principle fiber orientation, anisotropy of the peak fiber, and number of detected fibers. Based on these properties alone, we generated probabilistic maps of the inner and outer segments for both the DSI and MSI samples (Fig.7 A,B). Qualitative comparison of these maps shows a reliable and highly similar pattern of segmentation between the two pallidal regions. This is particularly evident in regions where outer and inner segments are divided along a curve approximately situated on the internal medullarly lamina in both hemispheres (Fig.7 A,B, and coronal slices Fig.7 D-E).

**Fig. 7.**
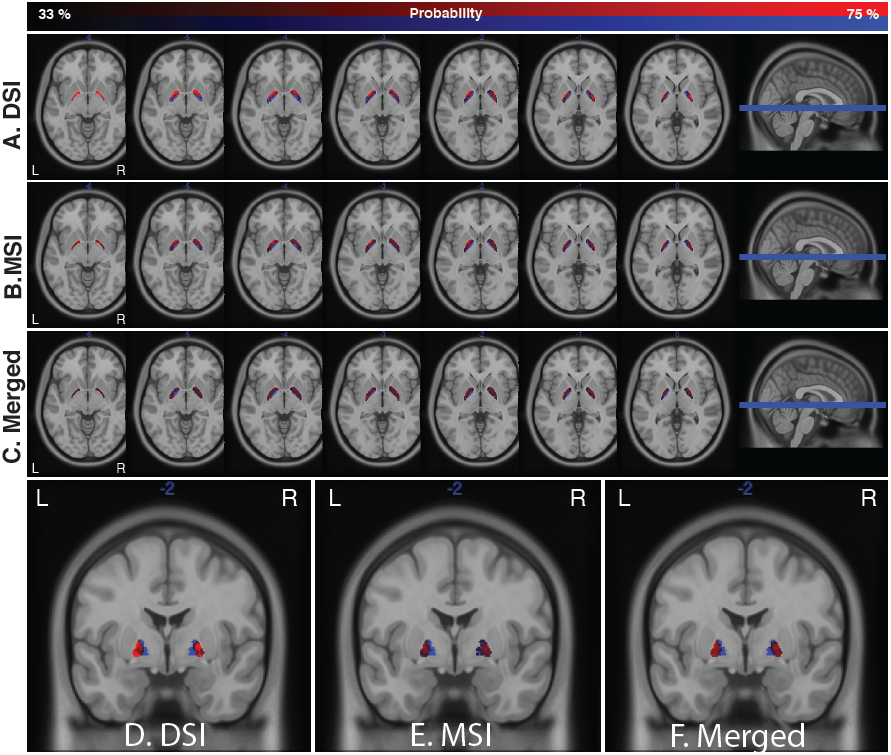
A-C. Probabilistic maps, across subjects, of the inner and outer segments in the DSI (A), MSI (B), and merged (C) datasets. Maps are thresholded between 33-77% probability. The background image in each image is the T1 ICBM template. Axial images span z coordinates [-6,0]. D-F. Coronal images of the same probabilistic maps at y = -2.

While these segmentation maps are not as clean as the hand-drawn maps based on the T1 signal (Fig. 2A), the general pattern of clustering is much better than expectations from chance. To explicitly quantify this, we compared the automatic segmentations to the hand drawn maps against chance accuracies generated from a permutation test (see Methods). Random accuracies ranged from 22% to 77% and were consistently higher in the inner segment (37% to 77%) relative to the outer segment (22% to 62%) reflecting the fact that there were fewer voxels within the inner segment and thus a higher chance of randomly overlapping with the correct assignment. As can be seen in Fig. 8, our classification significantly outperformed chance in all cases except the right hemisphere of the outer segment in the MSI dataset (Fig. 8C). Furthermore, accuracies were generally higher in the DSI sample than the MSI sample, likely due to the fact that the DSI sample was separable along all three features included in the clustering, while the MSI sample was not clearly separable based on the sensitivity curve measure (Fig. 6G-H).

**Fig. 8.**
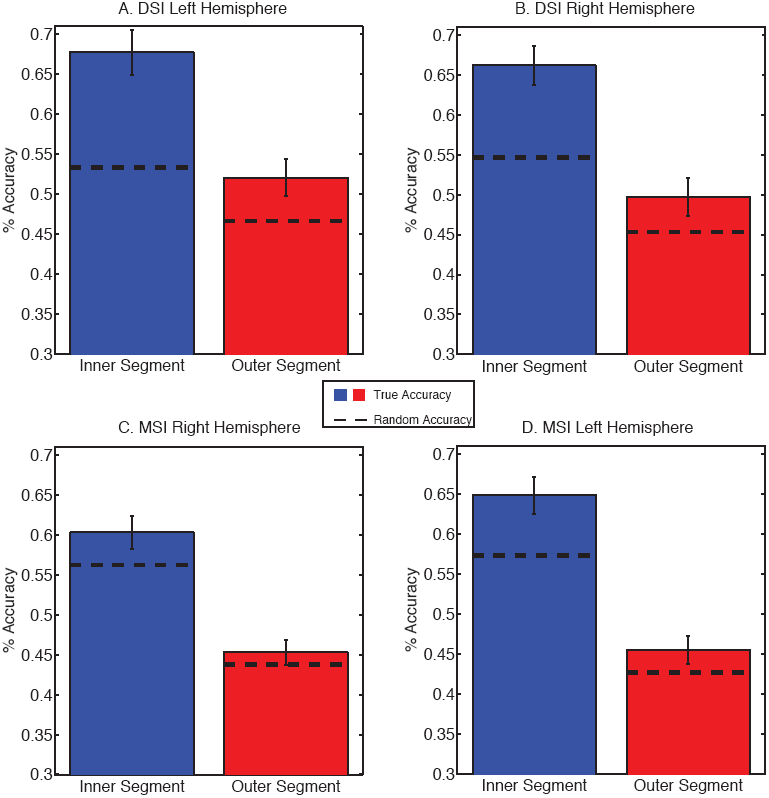
A-D. Mean accuracy results from k-means classification in the DSI (A,B) and MSI (C,D) datasets. Dotted lines show random accuracies obtained from 1000 iterations of a permutation test where clustered categories were scrambled. Error bars indicate 95% confidence intervals across subjects.

### Consistency across data sets

So far we have shown that both DSI and MSI samples exhibit similar differences in the pallidal segment diffusion signals and similar automatic parcellations of the internal and external pallidal masks. In order to quantify the similarity of the results between the two samples, we correlated the voxelwise probabilities between the DSI and MSI datasets (Fig. 9) for the internal and external segments separately. Overall, voxelwise probabilities between the two samples were moderately correlated in both hemispheres (r(138) =0.67 in left hemisphere vs. r(138) = 0.56 in the right hemisphere), suggesting that the SDF signal is capturing reliable topographic differences in underlying microstructural properties that is generally consistent across samples and the type of diffusion imaging approach used.

**Fig. 9.**
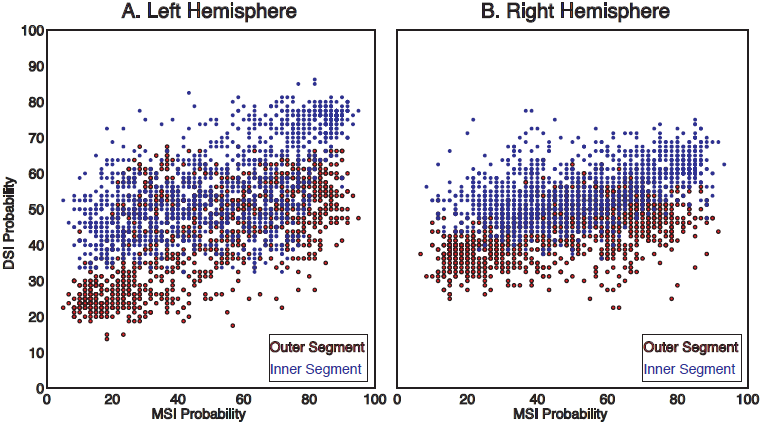
The voxelwise MSI probability (averaged across individual subjects) plotted against voxelwise DSI probability for each voxel in the left (A) and right (B) hemispheres. Each point corresponds to an individual voxel in either the outer (red) or inner (blue) segment.

Because both the DSI and MSI samples provided similar probability profiles, we aggregated both data sets to form a composite probabilistic map of the internal and external pallidal segments based on the underlying diffusion structure. These merged maps are shown in Figure 7C,F. Collapsing across the two acquisition methods revealed an even clearer distinction between the two pallidal nuclei. This confirms that classification-based purely on the properties of the diffusion signal is sufficiently robust across differences in acquisition approach and scan environments to capture the major divisions of the inner and outer segments of the globus pallidus.

## Discussion

For the first time we are able to show that the orientation distribution of the diffusion within the human pallidum is consistent with the presence of nigral and striatal efferents that run through these nuclei. Because a large portion of these pathways is buried within the pallidum, a region of high iron density, visualization of these efferents has been challenging with conventional imaging approaches. If diffusion MRI proves to be a reliable method of assessing the integrity of these pathways and their degradation in movement disorders, then quantifying degradation within the pallidum will be necessary to obtain accurate measurements of associated changes. Here we show that the SDF was also able to pick up on established histological differences between the internal and external segment of the globus pallidus (Hardman et al., 2002, Eid et al., 2013, Difiglia and Rafols 1988), resulting in the first automatic segmentation of these two nuclei. Thus, these measures are sufficiently robust to detect known differences in the pallidal segments. Furthermore, these differences in the diffusion signal between the internal and external segments were mostly consistent regardless of the acquisition method used (i.e., DSI vs. MSI) and able to classify the separate segments with accuracies well above chance expectations.

Although diffusion anisotropy measures are typically used to visualize pathways within core white matter regions of the brain, here we showed that tissue characteristics derived from the diffusion MRI signal, including differences in connectivity, intensity and sensitivity, can distinguish nuclear properties within the pallidum itself (see also Wiegell et al., 2003; Mang et al., 2012). We presume that orientations of the fibers in the two pallidal nuclei along with differences in density and myelination contribute to the characteristics of the SDFs within these voxels (Beaulieu, 2002). The predominately dorsolateral orientation of the resolved fiber peaks within the pallidal segments is consistent with the primary orientation of the striatopallidal, striatonigral and nigrostriatal/pallidal tracts (Wilson 1914, Szabo 1967, Szabo 1962, Fox and Rafols, 1976, Carpenter and Peter, 1972). The more pronounced dorsolateral orientation of the internal segment compared to the external segment in both the primary (Fig. 3) and secondary fibers (Fig. 4) is consistent with the volumetric differences between the two segments, since the nigrostriatal and striatonigral fibers traverse both segments. The medial shift observed in the outer segment (Fig. 3) is likely reflecting a distinct fiber system, possibly projections from the subthalamic nucleus. This open question can be resolved by a direct comparison of SDFs with postmortem histological analysis, which should be a goal of future work.

We should point out that the orientation of the peak fibers may not be completely consistent across diffusion imaging approaches. For example, there is a more pronounced medial shift in the external segment orientations in the DSI sample (Fig.3 A,B) than in the MSI sample (Fig.3 C,D). This may be due to the fact that the DSI sample is more sensitive to underlying microarchitectural features that contribute to a medial bias in the SDF signal. However, the secondary fiber in this sample was oriented in a more dorsal and dorsolateral direction (Fig. 4). This suggests that these nigral and striatal efferents are also present in the DSI sample, but to a weaker degree than in the MSI sample. This difference is likely due to the diffusion sampling scheme used in DSI and MSI. The DSI used a stronger diffusion sensitization strength (i.e. higher b-value) than the MSI, and it is more sensitive to restricted diffusion in gray matter.

Being able to detect signatures of the underlying cellular content of the pallidal nuclei *in vivo* has enormous potential as a biomarker for the integrity of basal ganglia pathways and their pathology. Many movement disorders involve degradation of specific pathways within the GPe and the GPi. In Parkinson’s disease for example, dopaminergic neurons within the substantia nigra degenerate which leads to a loss of the nigrostriatal fibers. Degradation of the nigrostriatal efferents in Parkinson’s patients have been recently identified using diffusion MRI (Ziegler et al., 2014). Given that the nigrostriatal fibers travel through the pallidum, degradation of these pathways may be reflected in the integrity of the microstructural architecture exhibited by the SDFs within the pallidum. Future comparative and clinical studies are needed in order to validate this assumption.

Beyond clinical implications, our results also have relevance to the investigation of basal ganglia function in neurologically healthy individuals. In the canonical direct-indirect pathway model of motor facilitation (Albin et al., 1989, DeLong 2000), activity in the GPe is correlated with inhibiting movement initiation, through disinhibition of the sub-thalamic nucleus, which in turn excites the GPi/SNr. Conversely, during movement facilitation, activity within the GPi decreases. Dysfunction of the direct and indirect pathways results in an imbalance between the two circuits, which causes impaired motor production as seen in Parkinson’s and Huntington’s disease. If the efficiency of processing with striatopallidal pathways is reflected in their microstructural integrity, then individual variation in performance on tasks may be correlated with the QA distributions of the striatopallidal fiber systems.

While our present results show promise for using the diffusion imaging signal as a measure of cellular architecture within sub-cortical nuclei, this approach still has some inherent limitations. First, as mentioned previously, diffusion imaging provides an indirect measure of cellular architecture. While validation work in non-human animals has provided insights into the underlying cellular properties for white matter using tensor-based reconstruction approaches (Wang et al., 2011, Wang et al., 2014, Thomas et al., 2014), model-based approaches have not been validated against histological models, particularly in gray matter (Blumenfeld-Katzir et al., 2011). Therefore, we do not know for sure what properties of the SDF reflect what properties of the underlying tissue. Future studies could probe precisely how changes in SDF properties are associated with variations in density, number of fibers, and myelination, by combining histological analysis and diffusion imaging in animal models and post mortem tissue analysis of the pallidum.

In addition, although we demonstrated that the boundaries of the pallidal nuclei are resolvable based solely on diffusion information, the segmentations are imperfect. In particular, there is a cluster of voxels in the anterior region of the pallidum that was misclassified in a significant number of subjects, and the parcellations were less accurate in the MSI data set as a whole. The proximity of the globus pallidus to the major white matter tracts of the internal capsule may contribute to partial voluming problems that contaminate the SDF signal in these voxels, resulting in classification errors. Future work could adaptively cluster using more sophisticated approaches to allow for noise clusters that could arise from errors in masking.

Despite this partial voluming problem, the segmentation results reported here still provide evidence of robust differences between the segments of the human pallidum. For example, unlike most subcortical parcellations (e.g. Behrens et al., 2003), we are not supplementing the clustering features with additional distance information that adds a strong prior on expected location of the nuclei. Such spatial priors would dramatically clean up the underlying maps; however, the distance from the expected nuclear location would become the dominant clustering feature. Although omitting these priors may lead to noisier segmentations, our approach provides a more robust measure for future studies to assess the pallidal cellular integrity in clinical populations.

Regardless of these limitations we have shown that the inner and outer segments of the globus pallidus not only express common asymmetries in their underlying SDFs, consistent with major efferent pathways, but also reliably differ among several properties of the diffusion signals. This was reliable enough that a simple and automatic clustering approach, based on properties of the SDF, resolved the inner and outer segments better than chance, regardless of the imaging acquisition used (i.e., DSI or MSI). This population atlas based analysis approach enables future studies to quantify the extent to which microstructural variability correlates with functional properties of the system, such as individual differences in inhibitory control ability and or clinical pathologies of the underlying fiber systems, providing a powerful new tool for investigating the cellular architecture of basal ganglia systems in vivo.

## Materials and Methods

### Participants and Acquisition

Two separate types of diffusion imaging were used for our analysis.

#### Diffusion Spectrum Imaging (DSI: CMU-60 Dataset)

Twenty nine male and thirty one female subjects were recruited from the local Pittsburgh community and the Army Research Laboratory in Aberdeen Maryland. All subjects were neurologically healthy, with no history of either head trauma or neurological or psychiatric illness. Subject ages ranged from 18 to 45 years of age at the time of scanning, with a mean age of 26 years (+/- 6 standard deviation). Six subjects were left handed (3 males, 3 females).

All participants were scanned on a Siemen’s Verio 3T system in the Scientific Imaging & Brain Research (SIBR) Center at Carnegie Mellon University using a 32-channel head coil. We collected a 50 min, 257-direction DSI scan using a twice-refocused spin-echo EPI sequence and multiple q values (TR = 9,916 ms, TE = 157 ms, voxel size = 2.4mm^3^, FoV = 231×231 mm, b-max = 5,000 s/mm^2^, 51 slices). Head-movement was minimized during the image acquisition through padding supports and all subjects were confirmed to have minimal head movement during the scan prior to inclusion in the template.

#### Multi-shell Imaging (MSI; HCP-80 Dataset)

The data were from the Human connectome project at WashU-Minnesota Consortium (Q1 release). Thirty six male and forty two female subjects were scanned on a customized Siemens 3T “Connectome Skyra” housed at Washington University in St. Louis. Subject ages ranged from 22-36 years of age at the time of scanning, with a mean age of 29.44 (+/- 3.5 standard deviation). All subjects were healthy, with no history of neurological or psychiatric illness. The two subjects that have subsequently been found by the HCP to exhibit gray matter heterotopia have been excluded from this analysis. The HCP DWI session was acquired using a spin-echo EPI sequence and (TR = 5520 ms, TE = 89.5 ms, voxel size = 1.25 mm^3^, FoV = 210×180, 3 shells of b = 1000, 2000, 3000 s/mm^2^, 111 slices, 90-directions for each shell).

### Diffusion MRI Reconstruction

All images were processed with a q-space diffeomorphic reconstruction method described previously (Yeh and Tseng, 2011) using DSI Studio (http://dsi-studio.labsolver.org/). The SDFs were reconstructed to a spatial resolution of 1 mm^3^. The white matter surface is rendered independently from an externally supplied 1 mm^3^ resolution white matter template. The quantitative anisotropy (QA; Yeh et al., 2010) and fiber orientation of the two major fibers in each voxel were exported into a separate file for analysis.

### SDF Analysis

Masks of the inner and outer segments of the pallidum were manually drawn by identifying the internal medullarly lamina in each hemisphere on the high resolution T1 ICBN MNI template. In each hemisphere, the GPe was drawn by including those voxels between the anterior and posterior limbs of the internal capsule, the putamen, and the internal medullary lamina (outer left/right in Fig.2A). Similarly, the GPi was drawn by including the voxels between the internal medullary lamina, the posterior limb and genu of the internal capsule (inner left/right in Fig.2A). All region of interest masks were drawn in MRICron (Rorden and Brett, 2000) and exported as NifTI images.

We then isolated the SDFs within each voxel of both region masks for analysis. For illustration, Fig. 2B shows a schematized version of a SDF illustrating three resolved fibers, with their independent magnitude (i.e., lengths, reflecting QA) and orientation. Two representative 3D SDFs from a voxel within the left GPi and a voxel within the left GPe are shown in Fig. 2C,D. For each voxel, we took three independent measures of the SDF structure: the principal fiber orientation, the number of resolved fibers across a range of QA thresholds, and QA magnitude of the principal fiber. To generate the angular distribution histograms (Figs. 3,4), we used the circstat toolbox (Berens 2009) and computed the circular mean of the voxel orientations across subjects. The internal capsule orientation (green arrows in Fig. 3,4) was calculated by averaging the orientations across a 4mm^3^ voxel cube situated prominently within the internal capsule in the left and right hemispheres. For plotting purposes, a gaussian smoothing kernel was applied to the QA maps (Fig. 6 A-D) for each subject (2 FWHM).

We extracted the primary fiber orientation, the number of fibers in each voxel, and the QA of the primary fiber from all voxels in each mask. Then we clustered the combined data across both masks using these diffusion features with a standard clustering approach in Matlab (R2014a). We specified two clusters, used squared euclidean distance as the distance metric and the k-means++ algorithm for cluster center initialization (replicates = 10). The clusters to the inner and outer segment were assigned based on the number of correctly assigned voxels relative to the hand drawn masks. This algorithm generated inner and outer segment maps for each subject and hemisphere in each dataset. To generate the probabilistic maps of each segment across all subjects in a sample (Fig. 7), voxel probabilities were estimated by averaging the binary categorization of each voxel in the inner and outer segment maps. Separate probability masks were calculated for in the DSI and MSI data sets, as well as across both samples (Merged). In order to quantify accuracy, we defined the classification accuracy as the number of voxels correctly assigned to the inner/outer segment from the k-means analysis using the manual segmentation as the correct assignment divided by the total number of voxels in that segment (Fig. 8).

Accuracy of the clustered segments was compared against the hand-segmented region of interest masks and a chance null distribution was estimated using a permutation procedure. On each iteration of the permutation test, every pallidal voxel was pseudorandomly assigned to either the inner or outer segment. The voxel’s permuted assignment was then compared to the voxel’s real assignment in the manually segmented pallidum masks and counted as correct if it matched that assignment. All of the correct assignments were counted for each iteration of the permutation test (n=1000). Chance accuracies were tallied across all iterations and the number of instances that the random assignment performed better than k-means classification was divided by the total number of iterations to generate the p-value for how well the automated classification performs against a classification based purely on chance.

## Acknowledgements

Data were provided [in part] by the Human Connectome Project, WU-Minn Consortium (Principal Investigators: David Van Essen and Kamil Ugurbil; 1U54MH091657) funded by the 16 NIH Institutes and Centers that support the NIH Blueprint for Neuroscience Research; and by the McDonnell Center for Systems Neuroscience at Washington University. This project was also supported by the Army Research Laboratory under Cooperative Agreement Number W911NF-10-2-0022. The views and conclusions contained in this document are those of the authors and should not be interpreted as representing the official policies, either expressed or implied, of the Army Research Laboratory or the U.S. Government. The U.S. Government is authorized to reproduce and distribute reprints for Government purposes notwithstanding any copyright notation herein. This research was also supported in part by T32 NS007433-17.

## References

Abhinav, K., Yeh, F.-C., Pathak, S., Friedlander, R. M., & Fernandez-Miranda, J. C. (2014). Advanced diffusion MRI fiber tracking in neurosurgical and neurodegenerative disorders and neuroanatomical studies: A review. Biochimica et Biophysica Acta (BBA) - Molecular Basis of Disease. doi:10.1016/j.bbadis.2014.08.002

Albin, R. L., Young, A. B., & Penney, J. B. (1989). The functional anatomy of basal ganglia disorders. Trends in Neurosciences, 12(10), 366–375. doi:10.1016/0166- 2236(89)90074-X

Alexander, G. E., DeLong, M. R., & Strick, P. L. (1986). Parallel organization of functionally segregated circuits linking basal ganglia and cortex. Annual Review of Neuroscience, 9, 357–381. doi:10.1146/annurev.neuro.9.1.357

Amunts, K., Lepage, C., Borgeat, L., Mohlberg, H., Dickscheid, T., Rousseau, M.-É.,… Evans, A. C. (2013). BigBrain: an ultrahigh-resolution 3D human brain model. Science (New York, N.Y.), 340(6139), 1472–5. doi:10.1126/science.1235381

Bastiani, M., Shah, N. J., Goebel, R., & Roebroeck, A. (2012). Human cortical connectome reconstruction from diffusion weighted MRI: The effect of tractography algorithm. NeuroImage, 62(3), 1732–1749. doi:10.1016/j.neuroimage.2012.06.002

Behrens, T. E. J., Johansen-Berg, H., Woolrich, M. W., Smith, S. M., Wheeler-Kingshott, C. A. M., Boulby, P. A.,… Matthews, P. M. (2003). Non-invasive mapping of connections between human thalamus and cortex using diffusion imaging. Nature Neuroscience, 6(7), 750–757. doi:10.1227/01.NEU.0000309595.77090.89

Beaulieu, C. (2002). The basis of anisotropic water diffusion in the nervous system - A technical review. NMR in Biomedicine. doi:10.1002/nbm.782

Berens, P. (2009). CircStat: A MATLAB toolbox for circular statistics. Journal of Statistical Software, 31, 1–21. doi:10.1002/wics.10

Beaulieu C. The basis of anisotropic water diffusion in the nervous syste. NMR Biomed. 2002; 15: 435–455

Blumenfeld-Katzir, T., Pasternak, O., Dagan, M., & Assaf, Y. (2011). Diffusion MRI of structural brain plasticity induced by a learning and memory task. PLoS ONE, 6(6). doi:10.1371/journal.pone.0020678

Burke, R. E., & O’Malley, K. (2013). Axon degeneration in Parkinson’s disease. Experimental Neurology. doi:10.1016/j.expneurol.2012.01.011

Carpenter, M. B., & Peter, P. (1972). Nigrostriatal and nigrothalamic fibers in the rhesus monkey. The Journal of Comparative Neurology, 144(1), 93–115. doi:10.1002/cne.901440105

Carpenter, M. B., & Mcmasters, R. E. (1964). Lesions of the Substantia Nigra in the Rhesus Monkey. Efferent Fiber Degeneration and Behavioral Observations. The American Journal of Anatomy, 114, 293–319.

Cossette, M., Lévesque, M., & Parent, A. (1999). Extrastriatal dopaminergic innervation of human basal ganglia. Neuroscience Research, 34(1), 51–54.

DeLong, M. R. (1990). Primate models of movement disorders of basal ganglia origin. Trends in Neurosciences, 13(7), 281–285. doi:10.1016/0166-2236(90)90110-V

Difiglia, M., & Rafols, J. A. (1988). Synaptic organization of the globus pallidus. Journal of Electron Microscopy Technique, 10(3), 247–263. doi:10.1002/jemt.1060100304

Eid, L., Champigny, M. F., Parent, A., & Parent, M. (2013). Quantitative and ultrastructural study of serotonin innervation of the globus pallidus in squirrel monkeys. European Journal of Neuroscience, 37(10), 1659–1668. doi:10.1111/ejn.12164

Fernández-Miranda, J. C., Wang, Y., Pathak, S., Stefaneau, L., Verstynen, T., & Yeh, F. C. (2014). Asymmetry, connectivity, and segmentation of the arcuate fascicle in the human brain. Brain Structure and Function. doi:10.1007/s00429-014-0751-7

Fox, C. A., & Rafols, J. A. (1976). The striatal efferents in the globus pallidus and in the substantia nigra. Research Publications - Association for Research in Nervous and Mental Disease, 55, 37–55.

Haber, S. N., Fudge, J. L., & McFarland, N. R. (2000). Striatonigrostriatal pathways in primates form an ascending spiral from the shell to the dorsolateral striatum. The Journal of Neuroscience!: The Official Journal of the Society for Neuroscience, 20(6), 2369–2382. doi:http://www.jneurosci.org/content/20/6/2369

Haber, S. N. (2003). The primate basal ganglia: Parallel and integrative networks. In Journal of Chemical Neuroanatomy (Vol. 26, pp. 317–330). doi:10.1016/j.jchemneu.2003.10.003

Hardman, C. D., Henderson, J. M., Finkelstein, D. I., Horne, M. K., Paxinos, G., & Halliday, G. M. (2002). Comparison of the basal ganglia in rats, marmosets, macaques, baboons, and humans: Volume and neuronal number for the output, internal relay, and striatal modulating nuclei. Journal of Comparative Neurology, 445(3), 238–255. doi:10.1002/cne.10165

Hollerman, J. R., Tremblay, L., & Schultz, W. (2000). Involvement of basal ganglia and orbitofrontal cortex in goal-directed behavior. In Progress in Brain Research (Vol. 126, pp. 193–215). doi:10.1016/S0079-6123(00)26015-9

Mang, S. C., Busza, A., Reiterer, S., Grodd, W., & Klose, A. U. (2012). Thalamus segmentation based on the local diffusion direction: A group study. Magnetic Resonance in Medicine, 67(1), 118–126. doi:10.1002/mrm.22996

Reiner, A., Albin, R. L., Anderson, K. D., D’Amato, C. J., Penney, J. B., & Young, A. B. (1988). Differential loss of striatal projection neurons in Huntington disease. Proceedings of the National Academy of Sciences of the United States of America, 85(15), 5733–5737. doi:10.1073/pnas.85.15.5733

Rorden, C., & Brett, M. (2000). Stereotaxic display of brain lesions. Behavioural Neurology, 12(4), 191–200.

Szabo, J. (1967). The efferent projections of the putamen in the monkey. Experimental Neurology, 19(4), 463–476. doi:10.1016/0014-4886(67)90166-5

Szabo, J. (1962). Topical distribution of the striatal efferents in the monkey. Experimental Neurology, 5(1), 21–36. doi:10.1016/0014-4886(62)90067-5

Thomas, C., Ye, F. Q., Irfanoglu, M. O., Modi, P., Saleem, K. S., Leopold, D. A., & Pierpaoli, C. (2014). Anatomical accuracy of brain connections derived from diffusion MRI tractography is inherently limited. Proceedings of the National Academy of Sciences, 111(46), 16574–16579. doi:10.1073/pnas.1405672111

Wang, Y., Fernández-Miranda, J. C., Verstynen, T., Pathak, S., Schneider, W., & Yeh, F. C. (2013). Rethinking the role of the middle longitudinal fascicle in language and auditory pathways. Cerebral Cortex, 23(10), 2347–2356. doi:10.1093/cercor/bhs225

Wang, Y., Wang, Q., Haldar, J. P., Yeh, F. C., Xie, M., Sun, P.,… Song, S. K. (2011). Quantification of increased cellularity during inflammatory demyelination. In Brain (Vol. 134, pp. 3587–3598). doi:10.1093/brain/awr307

Wang, X., Cusick, M. F., Wang, Y., Sun, P., Libbey, J. E., Trinkaus, K.,… Song, S. K. (2014). Diffusion basis spectrum imaging detects and distinguishes coexisting subclinical inflammation, demyelination and axonal injury in experimental autoimmune encephalomyelitis mice. NMR in Biomedicine, 27(7), 843–852. doi:10.1002/nbm.3129

Wedeen, V. J., Wang, R. P., Schmahmann, J. D., Benner, T., Tseng, W. Y. I., Dai, G.,… de Crespigny, A. J. (2008). Diffusion spectrum magnetic resonance imaging (DSI) tractography of crossing fibers. NeuroImage, 41(4), 1267–1277. doi:10.1016/j.neuroimage.2008.03.036

Wiegell, M. R., Tuch, D. S., Larsson, H. B., & Wedeen, V. J. (2003). Automatic segmentation of thalamic nuclei from diffusion tensor magnetic resonance imaging. Neuroimage, 19(2 Pt 1), 391–401. doi:10.1016/S1053-8119(03)00044-2

Wilson, S. a K. (1914). An experimental research into the anatomy and physiology of the corpus striatum. Brain, 36, 427–492.

Yeh, F. C., & Tseng, W. Y. I. (2011). NTU-90: A high angular resolution brain atlas constructed by q-space diffeomorphic reconstruction. NeuroImage, 58(1), 91–99. doi:10.1016/j.neuroimage.2011.06.021

Yeh, F. C., Wedeen, V. J., & Tseng, W. Y. I. (2010). Generalized q-sampling imaging. IEEE Transactions on Medical Imaging, 29(9), 1626–1635. doi:10.1109/TMI.2010.2045126

Ziegler, E., Rouillard, M., André, E., Coolen, T., Stender, J., Balteau, E.,… Garraux, G. (2014). Mapping track density changes in nigrostriatal and extranigral pathways in Parkinson’s disease. NeuroImage, 99, 498–508. doi:10.1016/j.neuroimage.2014.06.033

